# The Epstein-Barr virus ubiquitin deconjugase BPLF1 regulates the activity of Topoisomerase II during virus replication

**DOI:** 10.1101/2021.02.26.433008

**Authors:** Jinlin Li, Noemi Nagy, Jiangnan Liu, Soham Gupta, Teresa Frisan, Thomas Hennig, Donald P. Cameron, Laura Baranello, Maria G. Masucci

## Abstract

Topoisomerases are essential for the replication of herpesviruses but the mechanisms by which the viruses hijack the cellular enzymes are largely unknown. We found that topoisomerase-II (TOP2) is a substrate of the Epstein-Barr virus (EBV) ubiquitin deconjugase BPLF1. BPLF1 selectively inhibited the ubiquitination of TOP2 following treatment with topoisomerase poisons, interacted with TOP2α and TOP2β in co-immunoprecipitation and *in vitro* pull-down, stabilized Etoposide-trapped TOP2 cleavage complexes (TOP2cc) and promoted TOP2 SUMOylation, which halted the DNA-damage response and reduced Etoposide toxicity. Induction of the productive virus cycle promoted the accumulation of TOP2βcc, enhanced TOP2β SUMOylation, and reduced Etoposide toxicity in lymphoblastoid cell lines carrying recombinant EBV encoding the active enzyme. Attenuation of this phenotype upon expression of a catalytic mutant BPLF1-C61A impaired viral DNA synthesis and virus release. These findings highlight a previously unrecognized function of BPLF1 in promoting non-proteolytic pathways for TOP2cc debulking that favor cell survival and virus production.

## Introduction

Epstein–Barr virus (EBV) is a human gamma-herpesvirus that establishes life-long persistent infections in most adults worldwide. The virus has been implicated in the pathogenesis of a broad spectrum of diseases ranging from infectious mononucleosis (IM) to a variety of lymphoid and epithelial cell malignancies including both Hodgkin and non-Hodgkin lymphomas, undifferentiated nasopharyngeal carcinoma, and gastric cancer(Shannon-Lowe & Rickinson, 2019).

Like other herpesviruses, EBV establishes latent or productive infections in different cell types. In latency, few viral genes are expressed resulting in the production of proteins and non-coding RNAs that drive virus persistence and cell proliferation(Babcock et al., 1998). In contrast, productive infection requires the coordinated expression of a large number of immediate early, early and late viral genes, which leads to the assembly of progeny virus and death of the infected cells(Hammerschmidt & Sugden, 2013). Although much of the EBV-induced pathology has been attributed to viral latency, the importance of lytic products in the induction of chronic inflammation and malignant transformation is increasingly recognized(Munz, 2019), pointing to inhibition of virus replication as a useful strategy for preventing EBV associated diseases.

EBV replication is triggered by the expression of immediate early genes, which transcriptionally activates a variety of viral and host cell factors required for subsequent phases of the productive cycle(Countryman & Miller, 1985; Feederle et al., 2000; Murata, 2014). Among the cellular factors, DNA topoisomerase-I and -II (TOP1 and TOP2) were shown to be essential for herpesvirus DNA replication(Hammarsten et al., 1996; M Kawanishi, 1993; Wang et al., 2008), raising the possibility that topoisomerase inhibitors may serve as antivirals. Indeed, non-toxic concentrations of TOP1 and TOP2 inhibitors were shown to suppress EBV-DNA replication(M Kawanishi, 1993), and different TOP1 inhibitors reduced the transcriptional activity of the EBV immediate-early protein BZLF1 and the assembly of viral replication complexes(Wang et al., 2009). However, the mechanisms by which the virus harnesses the activity of these essential cellular enzymes remain largely unknown.

Topoisomerases sustain DNA replication, recombination and transcription by inducing transient single or double-strand DNA breaks that allow the resolution of topological problems arising from strand separation(Champoux, 2001; Wang, 2002). TOP2 homodimers mediate DNA disentanglement by inducing transient double strand-breaks (DSBs) through the formation of enzyme-DNA adducts, known as TOP2 cleavage complexes (TOP2ccs), between catalytic tyrosine residues and the 5’ends of the DSBs(Nitiss, 2009). Following the passage of the second DNA strand, TOP2 rejoins the DNA ends via reversion of the trans-esterification reaction. While TOP2-indued DSBs are relatively frequent in genomic DNA(Morimoto et al., 2019), failure to resolve TOP2ccs, as may occur upon endogenous or chemical stress that inhibits TOP2 activity, results in the formation of stable TOP2-DNA adducts that hinder DNA replication and transcription and trigger apoptotic cell death(Kaufmann, 1998). Thus, cellular defense mechanisms attempt to resolve the TOP2ccs via proteolytic or non-proteolytic mechanisms(Sun, Saha, et al., 2020). These may involve the displacement of TOP2 via ubiquitin(Mao et al., 2001) or SUMO and ubiquitin-dependent(Sun, Miller Jenkins, et al., 2020) proteasomal degradation, which, following the removal of residual peptide-DNA adducts by the Tyrosyl-DNA phosphodiesterase-2 (TDP2) resolving enzyme(Gao et al., 2014; Pommier et al., 2014), unmasks the DNA breaks and promotes activation of the DNA damage response (DDR)(Pommier et al., 2014). Alternatively, SUMOylation may induce conformational changes in the TOP2 dimer the expose the covalent bonds to the direct action of TDP2(Schellenberg et al., 2017). Two TOP2 isozymes expressed in mammalian cells share ∼70% sequence identity and have similar catalytic activities and structural features but are differentially regulated and play distinct roles in biological processes(Nitiss, 2009). While TOP2α is preferentially expressed in dividing cells and is essential for decatenating intertwined sister chromatids during mitosis(Chen et al., 2015), TOP2β is the only topoisomerase expressed in non-proliferating cells and is indispensable for transcription(Madabhushi, 2018), (McKinnon, 2016).

Ubiquitin-specific proteases, or deubiquitinating enzymes (DUBs), regulate protein turnover by disassembling poly-ubiquitin chains that target the substrate for proteasomal degradation(Komander, 2009). Several human and animal viruses encode DUB homologs that play important roles in the virus life cycle by promoting viral genome replication and inhibiting the host antiviral response(Bailey-Elkin et al., 2017; Gastaldello et al., 2010; Kattenhorn et al., 2005). In this study, we report that TOP2 interacts with and is a substrate of the DUB encoded in the N-terminal domain of the EBV large tegument protein BPLF1 and provide evidence for the capacity of BPLF1 to promote non-proteolytic pathways for the resolution of TOP2ccs, which enhances cell survival and virus replication.

## Results

### BPLF1 selectively inhibits the degradation of TOP2 in cells treated with topoisomerase poisons

To investigate whether the EBV encoded DUB regulates the proteasomal degradation of topoisomerases, FLAG-tagged versions of the 325 amino acid long N-terminal catalytic domain of BPLF1 and an inactive mutant where the catalytic Cys61 was substituted with Ala (BPLF1-C61A) were stably expressed by lentivirus transduction in HEK-293T cells under the control of a Tet-on regulated promoter (HEK-rtTA-BPLF1/BPLF1-C61A cell lines). Inducible expression was monitored by probing immunoblots of cells treated for 24 h with increasing concentration of doxycycline (Dox) with antibodies to the FLAG or V5 tags (Fig. S1A). Although the steady-state levels of BPLF1-C61A appeared to be lower, which may be due to rapid turnover, both polypeptides were readily detected by anti-FLAG immunofluorescence in approximately 50% of the induced cells, while the fluorescence was weak or below detection in the remaining cells (Fig. S1B).

To monitor ubiquitin-dependent proteasomal degradation, HEK-rtTA-BPLF1/BPLF1-C61A cells cultured overnight in the presence or absence of Dox were treated with the TOP1 poison Campthothecine or the TOP2 poison Etoposide in the presence or absence of the proteasome inhibitor MG132, and topoisomerase levels were assessed by western blot. Campthothecine and Etoposide trap TOP1-DNA and TOP2-DNA covalent adducts, respectively(Pommier, 2013), while MG132 prevents the proteasomal degradation of stalled topoisomerase-DNA intermediates(Mao et al., 2001). As expected, TOP1 was efficiently degraded in Campthothecine treated cells (Fig. 1A and Fig. 1C upper panels), while treatment with Etoposide promoted the degradation of both TOP2α and TOP2β (Fig. 1B and 1C middle and lower panels). The degradation was inhibited by treatment with MG132, confirming the involvement of the proteasome in the clearance of poisoned topoisomerases. Expression of wild type or mutant BPLF1 did not affect the Campthothecine-induced degradation of TOP1. In contrast, expression of BPLF1 was accompanied by stabilization of both TOP2α and TPO2β in Etoposide-treated cells, while the mutant BPLF1-C61A had no effect. Thus, the viral DUB selectively inhibits the degradation of TOP2 isozymes by the proteasome.

**Figure 1.**
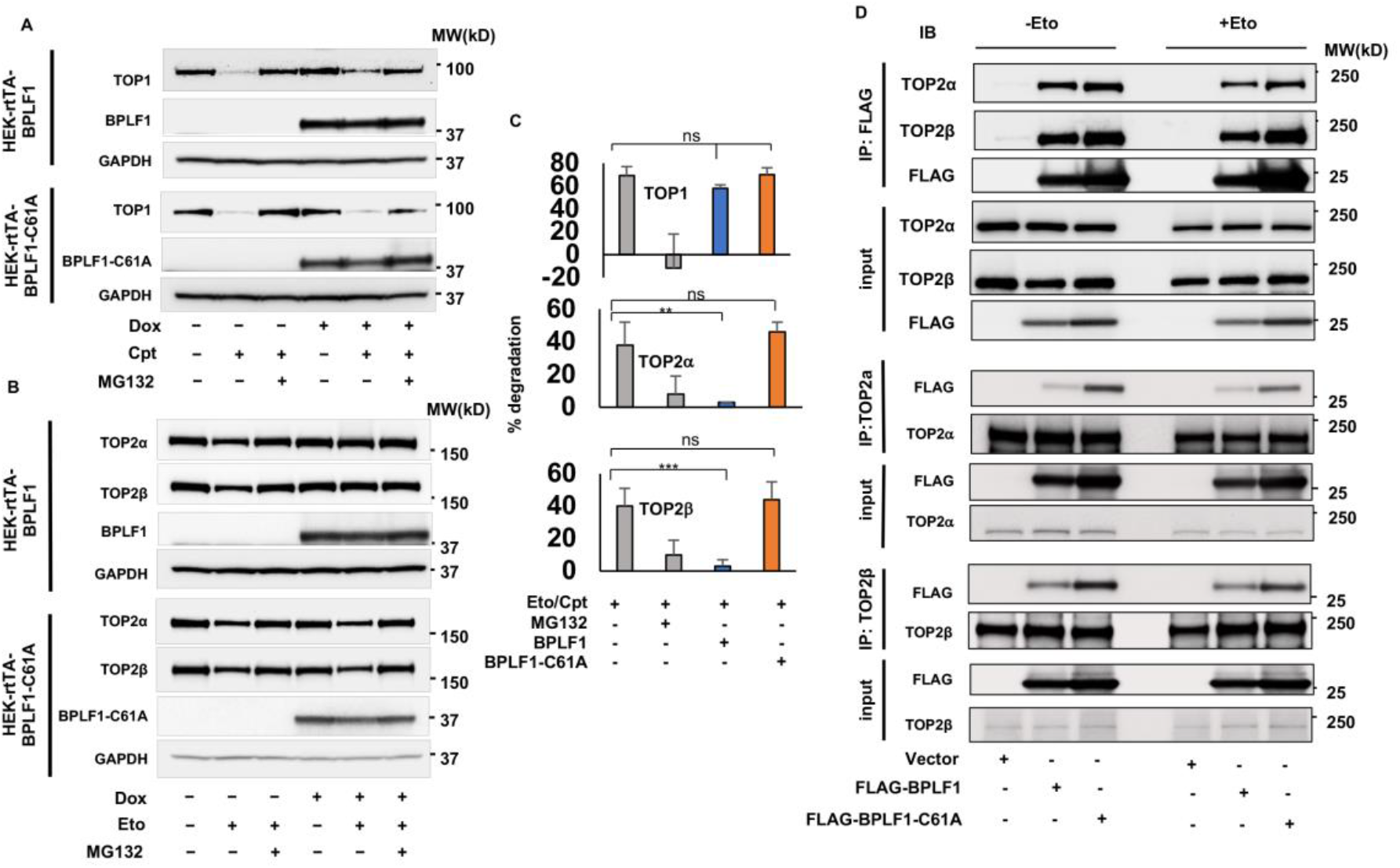
BPLF1 selectively binds to TOP2 and inhibits the degradation of TOP2 in cells treated with topoisomerase poisons. HEK-293T cell expressing inducible FLAG-BPLF1 or FLAG-BPLF1-C61A were seeded into 6 well plates and treated with 1.5 μg/ml doxycycline (Dox) for 24 h. After treatment for 3 h with 5 μM of the TOP1 poison Campthothecine (Cpt) or 6 h with 40 μM of the TOP2 poison Etoposide (Eto) with or without the addition of 10 μM MG132, protein expression was analyzed in western blots probed with the indicated antibodies. GAPDH was used as the loading control. (**A)** Representative western blots illustrating the expression of TOP1 in control and Campthothecine treated cells. The treatment induced degradation of TOP1 by the proteasome, which was not affected by the expression of BPLF1 or BPLF1-C61A in Dox treated cells. (**B**) Representative western blots illustrating the expression of TOP2α and TOP2β in Etoposide treated cells. Expression of BPLF1 reduced the Etoposide-induced degradation of both TOP2α and TOP2β while BPLF1-C61A had no appreciable effect. (**C**) The percentage degradation of TOP1, TOP2α and TOP2β in Campthothecine or Etoposide treated cells versus untreated controls was calculated from the intensity of the specific bands recorded in two (TOP1) or three (TOP2α and TOP2β) independent experiments using the ImageJ software. Data from HEK-rtTA-BPLF1/BPLF1-C61A cultured in the absence of Dox were pulled. Statistical analysis was performed using Student’s t-test. **P< 0.01; ***P < 0.001; ns, not significant. (**D**) HEK293T cells transfected with FLAG-BPLF1, FLAG-BPLF1-C61A, or FLAG-empty vector were treated with 40 μM Etoposide for 30 min and cell lysates were either immunoprecipitated with anti-FLAG conjugated agarose beads or incubated for 3 h with anti-TOP2α or TOPO2β antibodies followed by the capture of immunocomplexes with protein-G coated beads. Catalytically active and inactive BPLF1 binds to both TOP2α and TOP2β in both untreated and Etoposide treated cells (upper panels). Conversely, TOP2α (middle panels) and TOP2β (lower panels) interacts with both catalytically active and inactive BPLF1. Representative western blots from one of two independent experiments where all conditions were tested in parallel are shown.

### TOP2 is a BPLF1 substrate

To assess whether topoisomerases are direct substrates of BPLF1, we first investigated whether they interact in cells and in pull-down assays performed with recombinant proteins. Lysates of HEK-293T cells transiently transfected with FLAG-BPLF1 or FLAG-BPLF-C61A were immunoprecipitated with antibodies recognizing FLAG, TOP1, TOP2α or TOP2β. In line with the failure to rescue Campthothecine-induced degradation, BPLF1 did not interact with TOP1 (Fig. S2A), whereas both TOP2α and TOP2β were readily detected in western blots of the FLAG immunoprecipitates and, conversely, BPLF1 was strongly enriched in the TOP2α and TOP2β immunoprecipitates indicating that the proteins interact in cells (Fig. 1D). Of note, co-immunoprecipitation was more efficient when BPLF1-C61A was the bite, suggesting that TOP2 may be a substrate of the viral enzyme. To gain insight on the nature of the interaction, equimolar concentration of yeast expressed FLAG-TOP2α, or a TOP2α mutant lacking the unique C-terminal domain that is not conserved in the TOP2β isozyme, FLAG-TOP2α-ΔCTD, were mixed with bacterially expressed His-BPLF1 and reciprocal pull-downs were performed with anti-FLAG (Fig. S2B) or Ni-NTA coated beads (Fig. S2C). A weak BPLF1 band was reproducibly detected in western blots of the FLAG-TOP2α pull-downs probed with a His-specific antibody and, conversely, a weak FLAG-TOP2α band was detected in the His pull-downs. The binding of BPLF1 to TOP2α was not affected by deletion of the TOP2α C-terminal domain (Fig. S2D), pointing to a TOP2α and TOP2β shared domain shared as the likely site of interaction. Notably, comparison of the efficiency of *in vitro* pull-down versus co-immunoprecipitation suggests that binding may be stabilized by factors or TOP2 modifications that are only present in cell lysates. To further investigate whether the viral DUB deubiquitinates TOP2, BPLF1, TOP2α and TOP2β were immunoprecipitated from lysates of control and Etoposide-treated HEK-293T cells transiently transfected with BPLF1 or BPLF1-C61A and western blots were probed with a ubiquitin-specific antibody. The cell lysates were prepared under denaturing conditions to exclude non-covalent protein interactions and working concentrations of NEM and iodoacetamide were added to all buffers to inhibit DUB activity. In line with the capacity of Etoposide to promote the proteasomal degradation of TOP2, smears of high molecular weight species corresponding to ubiquitinated TOP2α and TOP2β were detected in the immunoprecipitates of Etoposide-treated cells (Fig. 2A). The intensity of the smears was strongly decreased in cells expressing catalytically active BPLF1, while the mutant BPLF1-C61A had no appreciable effect, confirming that TOP2 is a bona-fide BPLF1 substrate.

**Figure 2.**
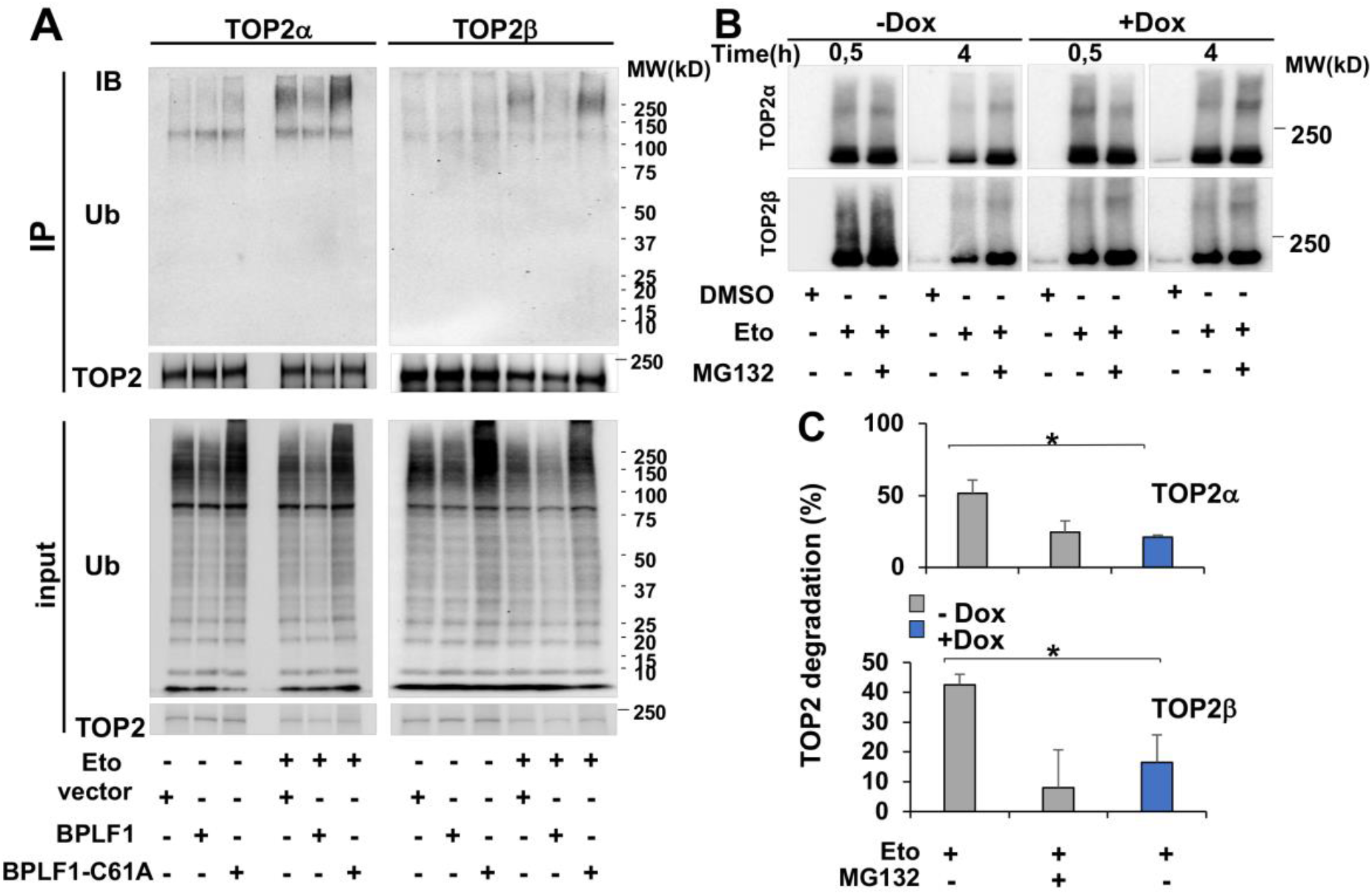
BPLF1 deubiquitinates TOP2 and stabilizes TOP2cc. **(A)** HEK293T cells were transiently transfected with plasmids expressing FLAG-BPLF1, FLAG-BPLF1-C61A, or the empty FLAG vector, and aliquots were treated with 40 μM Etoposide for 30 min. TOP2α and TOP2β were immunoprecipitated from cell lysates prepared under denaturing conditions in the presence of DUB inhibitors and western blots were probed with antibodies to TOP2α, TOP2β and ubiquitin. The expression of catalytically active BPLF1 inhibits the ubiquitination of TOP2α and TOP2β in Etoposide treated cells. Western blots from one representative experiment out of three are shown in the figure. **(B)** HEK-rtTA-BPLF1 cells were treated with 1.5 μg/ml Dox for 24 h followed by treatment with 80 μM Etoposide for the indicated time with or without the addition of 10 μM MG132. RADAR assays were performed as described in Materials and Methods and TOP2 trapped in 10 μg DNA was detected in western blots using antibodies to TOP2α or TOP2β. Trapped TOP2 appeared as a major band of the expected size and a smear of higher molecular weight species. The intensity of the trapped TOP2α and TOP2β bands decrease over time in control untreated cells due to proteasomal degradation, while the decrease was significantly reduced upon expression of BPLF1 in Dox treated cells. Western blots from one representative experiment out of two are shown in the figure. **(C**) The percentage of Etoposide-induced TOP2 degradation was calculated from the intensity of the specific bands measured with the ImageJ software. MG132 prevented the degradation of TOP2α and TOP2β trapped into TOP2cc in control BPLF1 negative cells whereas TOP2 degradation was significantly reduced in BPLF1 expressing cells. The mean ± SD of two independent experiments is shown in the figure.

Degradation of TOP2 by the proteasome plays an important role in the debulking of persistent TOP2ccs generated by topoisomerase poisons. To investigate whether the viral DUB may interfere with this process, HEK-rtTA-BPLF1 cells cultured for 24 h in the presence or absence of Dox were treated for 30 min or 4 h with Etoposide with or without addition of MG132, and DNA-trapped TOP2 was detected by RADAR (rapid approach to DNA adduct recovery) assays(Anand et al., 2018). Neither TOP2α nor TOP2β were detected in control DMSO treated cells confirming that only covalently DNA-bound species are isolated by this method (Fig. 2B). In Etoposide treated cells, TOP2α and TOP2β appeared as major bands of the expected size and smears of high molecular weight species that are likely to correspond to different types of post-translational modifications. Comparable amounts of trapped TOP2α and TOP2β were detected in cells treated with Etoposide for 30 min, independently of BPLF1 expression or MG132 treatment, indicating that neither treatment, either alone or in combination, affects the formation of TOP2ccs. As expected, the intensity of the TOP2 band decreased after Etoposide treatment for 4 h, which was inhibited by MG132, confirming the involvement of proteasome-dependent degradation in the debulking of Etoposide-induced TOP2ccs. The degradation of both TOP2α and TOP2β was significantly decreased at the 4 h time point in Dox treated cells, resulting in levels of stabilization comparable to those achieved by treatment with MG132 (Fig. 2C). Thus, BPLF1 deubiquitinates and stabilizes TOP2 trapped in covalent DNA adducts. The finding was independently confirmed in experiments where TOP2ccs were stabilized by alkaline lysis(Ban et al., 2013) (Fig. S3A). Smears of high molecular weight species were readily detected above the main TOP2β band in Dox-induced Etoposide-treated HEK-rtTA-BPLF1 cells, whereas high molecular weight species were not detected when the blots were probed with a TOP1 specific antibody, confirming that the high molecular weight species correspond to DNA-trapped TOP2 (Fig. S3A). As expected, the intensity of the high molecular weight species decreased with time in BPLF1 negative cells, and the decrease was inhibited by MG132 confirming the involvement of proteasomal degradation. In cells expressing catalytically active BPLF1, the intensity of the high molecular weight species remained virtually constant over the observation time, resulting in significantly higher amounts of residual TOP2ccs (Fig. S3B). Similar results were obtained when the blots were probed with antibodies to TOP2α.

### BPLF1 inhibits the detection of Etoposide-induced DNA damage and promotes TOP2 SUMOylation and cell survival

The removal of TOP2 trapped in TOP2ccs induces a DNA damage response (DDR) that, while limiting Etoposide toxicity, may also promote genomic instability and apoptosis(Lee et al., 2016; Mao et al., 2001; Sciascia et al., 2020). To test whether the capacity of BPLF1 to stabilize TOP2ccs interferes with DDR activation, HEK-rtTA-BPLF1/BPLF-C61A cells cultured in the presence or absence of Dox for 24 h and then treatment with Etoposide for 1 h. The accumulation of phosphorylated histone H2AX (γ H2AX), a validated DDR marker(Mah et al., 2010), was monitored by immunofluorescence in BPLF1 positive and negative cells. As illustrated by representative fluorescence micrographs (Fig. 3A, upper panels) and plots of fluorescence intensity in BPLF1 positive and negative cells (Fig. 3B, upper panels), a diffuse γ H2AX fluorescence was readily detected in Etoposide-treated BPLF1 negative cells and in cells expressing the mutant BPLF-C61A.

**Figure 3.**
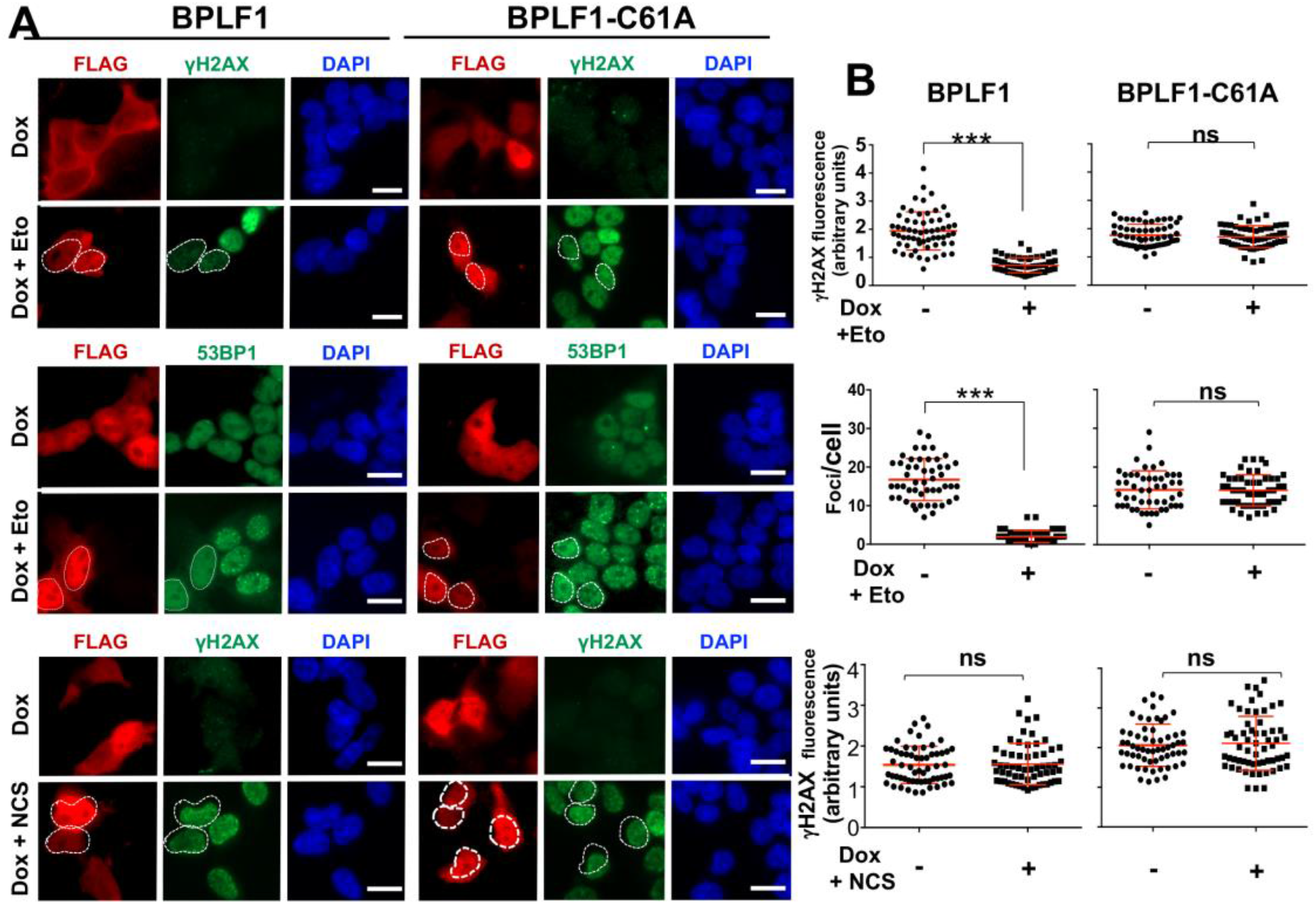
BPLF1 selectively inhibits the detection of TOP2-induced DNA damage. HEK-rtTA-BPLF1/BPLF1-C61A cells grown on cover-slides were treated with 1.5 μg/ml Dox for 24 h to induce the expression of BPLF1 followed by treatment for 1 h with 40 μ Etoposide or 0.5 μg/ml of the radiomimetic Neocarzinostatin (NCS) before staining with the indicated antibodies. (**A**) The cells were co-stained with antibodies against FLAG (red) and antibodies to γH2AX or 53BP1 (green) and the nuclei were stained with DAPI (blue). Expression of the catalytically active BPLF1 was associated with a significant decrease of nuclear γH2AX fluorescence and decreased formation of 53BP1 foci while the BPLF1-C61A mutant had no effect. Neither the catalytically active nor the inactive BPLF1 affected the induction of γH2AX in cells treated with NCS. Representative micrographs from one out of two experiments where all conditions were tested in parallel are shown. Scale bar = 10 μm. **(B)** Quantification of γH2AX fluorescence intensity and 53BP1 foci in BPLF1/BPLF1-C61A positive and negative cells from the same image. The Mean ± SD of fluorescence intensity in at least 50 BPLF1-positive and 50 BPLF1-negative cells recorded in each condition is shown. Statistical analysis was performed using Student’s t-test. ***P <0.001; ns, not significant.

Cells expressing active BPLF1 showed significantly weaker γH2AX fluorescence, suggesting that the viral enzyme counteracts DDR activation. Accordingly, DNA repair was not triggered as assessed by the impaired formation of 53BP1 foci in BPLF1 positive compared to negative cells (Fig. 3A and 3B, middle panels). A comparable BPLF1-dependent decrease of γH2AX fluorescence and formation of 53BP1 and BRCA1 foci was observed upon Etoposide treatment in HeLa cells transiently transfected with BPLF1/BPLF1-C61A (Fig. S4), confirming that the effect is not cell-type specific. To assess whether the failure to activate the DDR may be due to the capacity of BPLF1 to target events downstream of the formation of DSBs, cells expressing BPLF1/BPLF1-C61A were treated with the radiomimetic agent Neocarzinostain (NCS)(Povirk, 1996). Neither BPLF1 nor BPLF1-C61A altered the induction of γH2AX in NCS treated cells (Fig. 3A and 3B, lower panels). Thus, BPLF1 selectively inhibits the DDR and DNA repair responses triggered by Etoposide-induced DSBs.

In the absence of TOP2 degradation, TOP2cc may be resolved via a non-proteolytic process whereby SUMOylation-dependent conformational changes expose the tyrosyl-DNA bond to the activity of Tyrosyl-DNA-phosphodiesterase-2 (TDP2), which enables DSBs repair without the need of nuclease activity(Schellenberg et al., 2017). To assess whether this pathway may be engaged in BPLF1 expressing cells, Dox-treated HEK-rtTA-BPLF1 cells were exposed to Etoposide for 30 min and western blots of TOP2ccs isolated by RADAR were probed with antibodies to ubiquitin and SUMO2/3. As expected, smears of high molecular weight species reacting with both ubiquitin-and SUMO2/3-specific antibodies were highly enriched in Etoposide treated cells (Fig. 4A).

**Figure 4.**
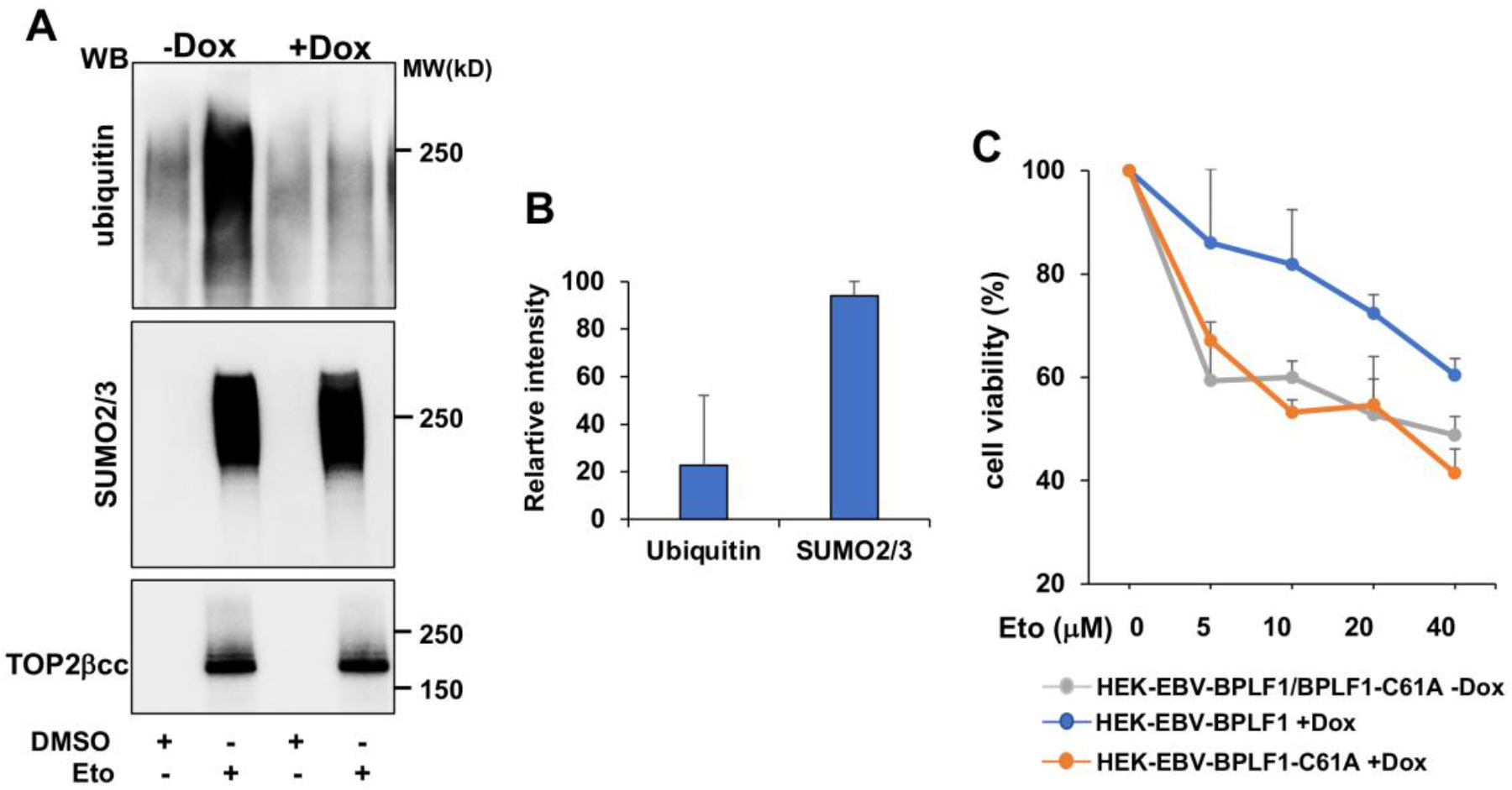
BPLF1 promotes TOP2 SUMOylation and cell viability following Etoposide treatment. **(A)** HEK-rtTA-BPLF1 cells were cultured for 24 h in the presence or absence of 1.5 μg/ml Dox and then treated with 80 μM Etoposide for 30 min followed by detection of DNA trapped TOP2 by RADAR assay. Western blots of proteins bound to 10 μg DNA were probed with antibodies to ubiquitin, SUMO2/3 and TOP2. The expression of BPLF1 was associated with strongly decreased ubiquitination of the TOP2ccs while SUMOylation was only marginally affected. **(B)** The intensity of the ubiquitin, SUMO2/3 and TOP2bcc specific bands was quantified by densitometry using the ImageJ software. Relative intensity was calculated as the % intensity in Dox-treated versus untreated cells after normalization to TOP2cc. Mean ± SD of two independent experiments. (**C**) HEK-rtTA-BPLF1/BPLF1-C61A cells were cultured for 24 h in the presence or absence of 1.5 μg/ml Dox and then treated overnight with the indicated concentration of Etoposide before assessing cell viability by MTT assays. The expression of catalytically active BPLF1 decreased the toxic effect of Etoposide over a wide range of concentrations while BPLF1-C61A had no appreciable effect. The mean ± SD of two independent experiments is shown.

Although the formation of TOP2ccs was not affected (Fig. 4A, lower panel), expression of the viral DUB was accompanied by a dramatic decrease of ubiquitinated species, while the intensity of the SUMO2/3 smear was largely unaffected (Fig 4A upper and middle panels and 4B). Thus, in addition to preventing the detection of TOP2-induced DNA damage, by inhibiting TOP2 ubiquitination BPLF1 may shift the processing of TOP2ccs towards non-proteolytic pathways, which could counteract the toxic effect of Etoposide. To test this possibility, cell viability was assessed by Thiazolyl blue tetrazolium bromide (MTT) assays in untreated and Dox-induced HEK-rtTA-BPLF1/BPLF1-C61A cells following overnight exposure to increasing concentrations of Etoposide. Similar levels of cell viability were observed in cells treated with Etoposide in the absence of Dox whereas, in line with the hypothesized protective effect of BPLF1, cell viability was consistently improved in cells expressing the active enzyme over a wide range of Etoposide concentrations and the BPLF1-C61A mutant had no appreciable effect (Fig. 4C).

### BPLF1 regulates the activity of TOP2β during productive infection

In the final set of experiments, we asked whether physiological levels of BPLF1 regulate the activity of TOP2 during the productive virus cycle in EBV infected cells. To this end, infectious virus rescued from HEK-293-EBV cells carrying recombinant EBV expressing wild type or mutant BPLF1(Gupta et al., 2018) was used to transform normal B-lymphocytes into immortalized lymphoblastoid cell lines (LCL-EBV-BPLF1/BPLF1-C61A). To optimize the induction of the productive virus cycle, the LCLs were stably transduced with a recombinant lentivirus expressing the viral transactivator BZLF1 under the control of a tetracycline-regulated promoter. Treatment with Dox induced the expression of early (BMRF1) and late (BFRF3) viral antigens detected in western blots probed with specific antibodies (Fig. S5A), and BPLF1 mRNA detected by qPCR (Fig. S5B). Of note, the induction of late antigens was consistently weaker in LCL-EBV-BPLF1-C61A (Fig. S5A), pointing to impairment of the late phase of the virus cycle.

Consistent with the establishment of a pseudo-S-phase where progression to G2/M is blocked and cellular DNA synthesis is inhibited(Kudoh et al., 2003), induction of the productive cycle was associated with a strong decrease of TOP2α mRNA (Fig. S5C) and protein levels (Fig. 5A) while TOP2β protein and mRNA showed either no change or a small increase of protein levels in cells expressing catalytically active BPLF1 (Fig. 5A and S5C).

**Figure 5.**
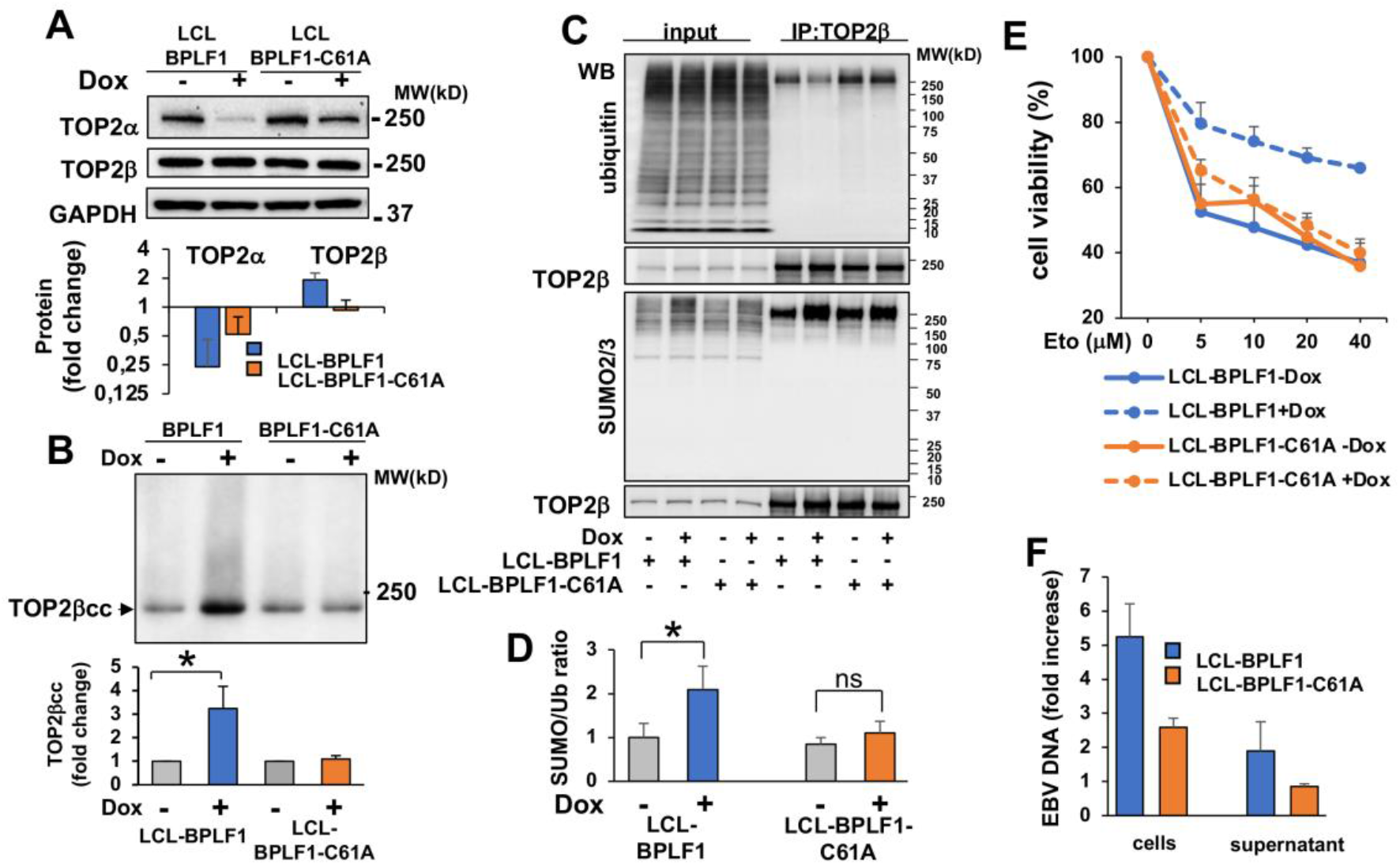
BPLF1 regulates the expression and activity of TOP2β during productive infection. The productive virus cycle was induced by treatment with 1.5 mg/ml Dox in LCL cells carrying recombinant EBV encoding wild type or catalytic mutant BPLF1 and expressing a tetracycline regulated BZLF1 transactivator. (**A**) The expression of TOP2α and TOP2β was assessed by western blot and the intensity of the specific bands was quantified using the ImageJ software. Induction of the productive cycle was associated with a highly reproducible downregulation of TOP2α while TOP2β was either unchanged or slightly increased. The effect was stronger in cells expressing wild type BPLF1. Representative western blots and quantification of specific bands in three to five independent experiments are shown. (**B)** The formation of TOP2bcc was investigated by RADAR assays in untreated and induced LCLs. Representative western blot illustrating the significant increase of TOP2bcc upon induction of the productive virus cycle in LCL cells expressing catalytically active BPLF1. BPLF1-C61A had no appreciable effect. One representative western blots and quantification of the intensity of the TOP2b smears in three independent experiments are shown. Fold increase was calculated as the ratio between the smear intensity in control versus induced cells. *P<0.05. (**C**) TOP2β was immunoprecipitated from total cell lysates of control and induced LCLs and western blots were probed with antibodies to TOP2β, ubiquitin and SUMO2/3. Western blots illustrating the decreased ubiquitination and increased SUMOylation of TOP2β in cells expressing catalytically active BPLF1. One representative experiment out of three is shown. (**D**) The intensity of the bands corresponding to immunoprecipitated TOP2β, ubiquitinated and SUMOylated species was quantified using the ImageJ software and the SUMO/Ub ratio was calculated after normalization to the intensity of immunoprecipitated TOP2b. The mean ± SD of three independent experiments is shown. *P<0.05. (**E**) The productive cycle was induced in LCL-EBV-BPLF1/BPLF1-C61A by culture for 72 h in the presence 1.5 μg/ml Dox. After washing and counting, 5×10^4^ live cells were seeded in triplicate wells of 96 well plates and treated overnight with the indicated concentration Etoposide before assessing cell viability by MTT assays. The expression of catalytically active BPLF1 enhanced cell viability over a wide range of Etoposide concentration with BPLF1-C61A had no appreciable effect. The mean ± SD of cell viability in three independent experiments is shown. **(F)** The amount of cell associated and release EBV DNA was measures in the cell pellets ad supernatants after induction for 72 h. Fold induction was calculated relative to uninduced cells. Mean ± DS of 3 experiments.

This was associated with a significant increase of TOP2βccs relative to uninduced cells or cells expressing the mutant BPLF1-C61A (Fig. 5B), and with decreased TOP2β ubiquitination (Fig. 5C top panel). As previously reported, higher molecular weight species detected by the SUMO2/3 specific antibody were increased in induced cells due to viral micro-RNA-dependent downregulation of RNF4(Li et al., 2017). SUMOylated species were also increased in immunoprecipitated TOP2β (Fig. 5C lower panel), resulting in a significant shift of the SUMO/ubiquitin ratio towards TOP2β SUMOylation in cells expressing catalytically active BPLF1 (Fig. 5D). As observed with the inducible HEK-rtTA-BPLF1 cell line, expression of the active viral DUB counteracted the toxic effect of Etoposide (Fig. 5E). Furthermore, the BPLF1-mediated regulation of TOP2β expression and ubiquitination was associated with higher levels of viral DNA replication and efficient release of infectious virus particles as measured by qPCR in cell pellets and culture supernatants (Fig. 5F).

## Discussion

Although compelling evidence points to a pivotal role of topoisomerases in the replication of herpesviruses and other DNA viruses(M. Kawanishi, 1993; Wang et al., 2009; Wang et al., 2008), very little is known about the mechanisms by which the viruses harness the activity of these cellular enzymes. In this study, we have shown that the ubiquitin deconjugases encoded in the N-terminal domain of the EBV large tegument protein BPLF1 regulates the activity of TOP2β during productive EBV infection by promoting the proteasome-independent debulking of TOP2-DNA adducts, which favors cell survival and the faithful replication and transcription of viral DNA. The findings highlight a previously unrecognized function of the viral enzyme in hijacking cellular functions that enable efficient virus production. Our proposed model for the activity of BPLF1 is shown in Fig. 6.

**Figure 6.**
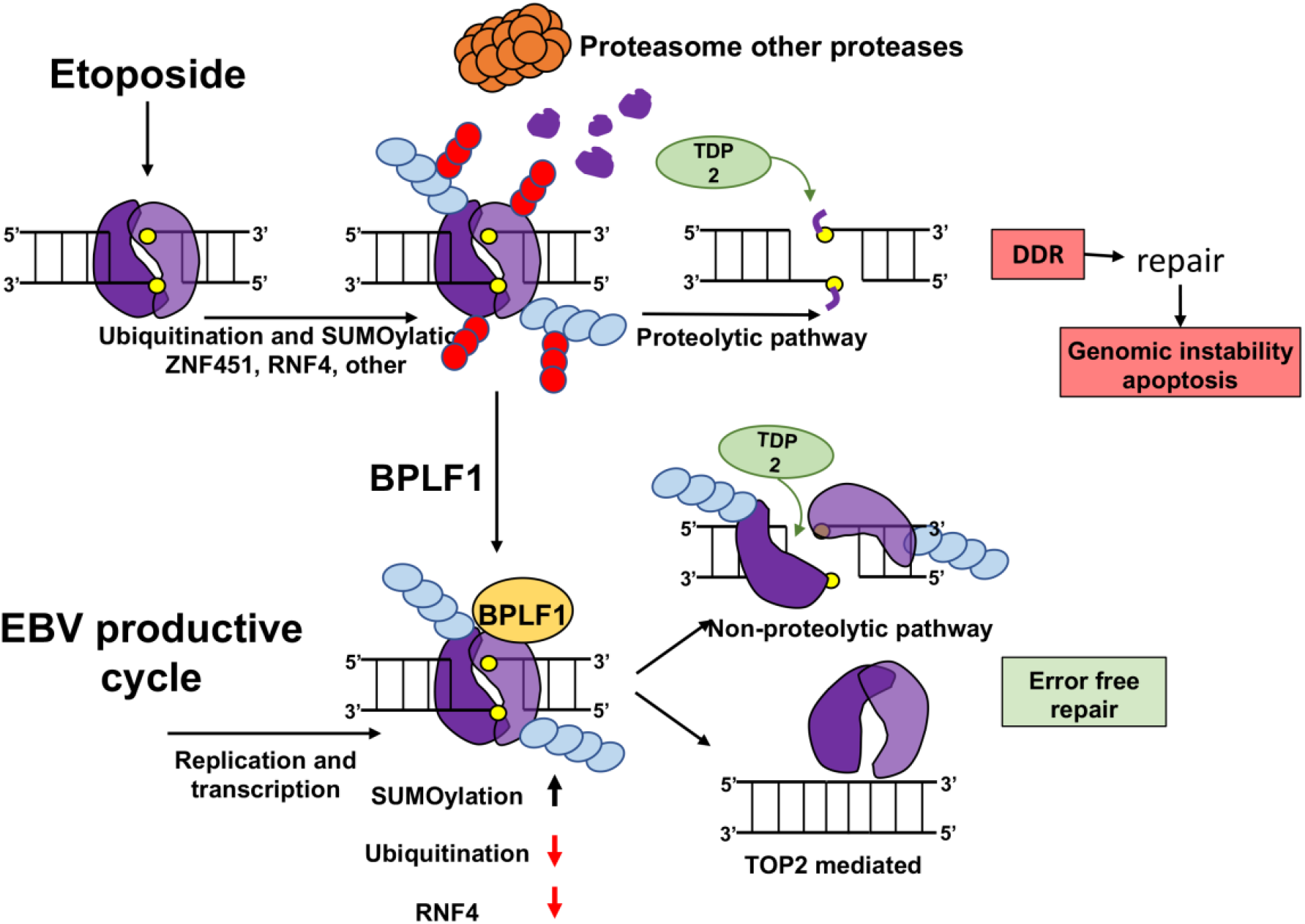
Model of TOP2 regulation by the BPLF1. TOP2 (violet) trapped in TOP2ccs (yellow) is targeted for proteasomal degradation via SUMOylation (light blue) and ubiquitination (red) mediated by the SUMO ligase ZNF451, the SUMO-targeting ubiquitin ligase RNF4 and other cellular ubiquitin ligases, leading to the display of partially digested 5’-phosphotyrosyl-DNA adducts. Processing by the TDP2 resolvase generates protein-free DSBs that trigger the DDR and error-free or error-prone DNA repair. Imprecise repair leads to apoptosis and genomic instability. BPLF1 inhibits the degradation of Etoposide-poisoned TOP2, which inhibits activation of the DDR. In the absence of proteasomal degradation, SUMOylation may alter the conformation of the TOP2 dimer allowing direct access of TDP2 to the 5’-phosphotyrosyl-DNA bonds, which promotes error-free repair. During productive EBV infection, the concomitant expression of BPLF1 and downregulation of RNF4 favors the accumulation of SUMOylated TOP2β and the activation of non-proteolytic pathways for TOP2ccs debulking, which, in the absence of TOP2 poisons, may be mediated by TOP2 itself. This ensures the fidelity of virus replication and transcription and enhances cell survival and virus production.

We found that the viral DUB that is physiologically released during productive infection via caspase-1-mediated cleavage of the EBV large tegument protein BPLF1(Gastaldello et al., 2013) selectively binds to TOP2α and TOP2β, and effectively counteracts their ubiquitination and proteasomal degradation in cells treated with Etoposide (Fig. 1, Fig 2A). In the absence of proteasomal degradation, TOP2ccs were stabilized (Fig 2B, 2C), which prevented the unmasking of TOP2-induced DSBs (Fig 3) and promoted resistance to the toxic effect of Etoposide (Fig 4C). Several lines of evidence suggest that the potent DDR triggered by the proteolytic debulking of TOP2ccs may be detrimental to cell survival and genomic integrity. Following the degradation of TOP2, protein-free DSBs engage multiple pathways for error-free or error-prone repair, including MRE11 nuclease-dependent homologous recombination (HR)(Hoa et al., 2016) and non-homologous end joining (NHRJ)(Gomez-Herreros et al., 2013; Gomez-Herreros et al., 2017). Recent findings suggest that a substantial fraction of the Etoposide induced DSBs undergo extensive DNA end-resection(Sciascia et al., 2020), which favors mispairing and the occurrence of chromosomal rearrangements that compromise cell viability or promote genomic instability. Of note, these genotoxic effects were efficiently counteracted by inhibition of the proteasome prior or during Etoposide treatment, supporting the notion that the non-proteolytic resolution of TOP2ccs can minimize DSB misrepair and promote genomic integrity(Sciascia et al., 2020).

We found that expression of the catalytically active viral deubiquitinase closely mimicked the stabilization of TOP2ccs (Fig 2B) and inhibition of both DDR activation (Fig. 3) and Etoposide toxicity (Fig. 4C and 5E) observed upon inhibition of the proteasome. While in line with the notion that ubiquitination is strictly required for the targeting of substrates to the proteasome, this finding points to the capacity of BPLF1 to shift the cellular strategy for TOP2cc debulking towards proteasome-independent pathways that may ensure higher fidelity of DNA repair and reduce toxicity. In this context, it is important to notice that, while inhibiting ubiquitination, catalytic active BPLF1 did not affect the SUMOylation of TOP2cc in Etoposide treated cells (Fig. 4A) and promoted the preferential SUMOylation of TOP2β during productive EBV infection (Fig. 5C and 5D). SUMOylation plays multiple roles in the debulking of TOP2ccs. It may mediate TOP2 proteolysis by serving as a recognition signal for ubiquitination mediated by SUMO-targeted ubiquitin ligases such as RNF4(Sun, Miller Jenkins, et al., 2020) or may recruit SprT-family metalloproteases, such as SPRTN(Lopez-Mosqueda et al., 2016) and ARC/GCNA(Borgermann et al., 2019), that are involved in the proteasome-independent proteolytic debulking of DNA-protein adducts. Interestingly, the activity of SPRTN is inhibited by mono-ubiquitination(Stingele et al., 2016) and yet unpublished findings suggest that failure to deubiquitinate SPRTN upon depletion of the cellular deubiquitinase USP11 leads to the accumulation of unrepaired DNA-protein adducts(Perry et al., 2020). Thus, BPLF1 could mimic the activity of the cellular DUB. In addition, SUMOylation of TOP2 by the ZNF451 ligase was shown to promote the non-proteolytic resolution of TOP2-DNA cross-links via direct recruitment of TDP2 through a “split-SIM” SUMO2 engagement platform(Schellenberg et al., 2017). SUMOylation was shown to alter the conformation of the trapped TOP2 dimers, thereby facilitating the access of TDP2 to the tyrosyl-DNA covalent bond and promoting error-free rejoining of the DSBs. This may be accomplished by the T4 DNA ligase(Gomez-Herreros et al., 2013) or, upon removal of Etoposide, by TOP2 (Sciascia et al., 2020).

While the pivotal role of topoisomerases in both the latent and lytic replication of herpesviruses is firmly established(Benson & Huang, 1988; Ebert et al., 1990; M Kawanishi, 1993), the roles of the individual enzymes are not well understood. The torsion-relieving activity of TOP1 was shown to be essential for reconstitution of the HSV replication machinery using purified viral proteins(Nimonkar & Boehmer, 2004), and its recruitment to the viral replication complex was required for the lytic origin (OriLyt)-driven replication of EBV(Wang et al., 2009) and KSHV(Wang et al., 2008). Less is known about the function of the TOP2 isozymes, although the importance of TOP2 is underscored by its upregulation during the productive cycle of HCMV(Benson & Huang, 1990) and KSHV(Wang et al., 2008) despite a general host proteins shutoff during virus replication. We have found that TOP2α mRNA is strongly downregulated upon induction of the productive virus cycle in EBV infected cells (Fig. S5C) and the protein becomes virtually undetectable in cells expressing catalytically active BPLF1 (Fig. 5A). While possibly related to the virus-induced arrest of the cell cycle in G1/S, the precise mechanism of this downregulation remains unknown. Nevertheless, our findings exclude a major role of TOP2α in the replication of the viral genome. In contrast, the expression of TOP2β was either not affected or slightly upregulated during productive infection, which is in line with the exclusive expression of this topoisomerase in resting cells and its essential role in transcription. Most importantly, we found that catalytically active BPLF1 was required for the accumulation of TOP2βccs in cell entering the productive cycle (Fig 5B), which, in the absence of topoisomerase poisons, is likely to indicate a significant increase of TOP2β activity driven by viral DNA replication and/or transcription. Conceivably, the capacity of BPLF1 to stabilize TOP2ccs and their resolution by non-proteolytic pathways that favor error-free repair and cell survival may be instrumental to ensure faithful and proficient replication and transcription of the viral genome. Of note, the BPLF1-mediated salvage of TOP2β from proteolytic disruption is likely to be reinforced by the concomitant downregulation of RNF4(Li et al., 2017), which may ensure that sufficient levels of the protein remain available throughout the productive cycle to sustain efficient virus production.

Aberrant expression of BPLF1 in the context of abortive lytic cycle reactivation has been reported in EBV associated malignancies such as undifferentiated nasopharyngeal carcinoma, NK-T cell lymphomas, and a subset of gastric cancers ^(Peng et al., 2019)^,^(Borozan et al., 2018)^. Etoposide and other topoisomerase poisons are used clinically as therapeutic anticancer agents against these malignancies(Delgado et al., 2018). Our data suggest that the expression of BPLF1 could be potentially used as a biomarker to predict the effectiveness of chemotherapeutic regimens that incorporate topoisomerase poison.

## Acknowledgements

We are deeply grateful to Dr Arne Lindquist and Prof Nico Dantuma (CMB, Karolinska Institutet) for generously sharing knowledge and reagents and to Ms. Lisa Wohlgemuth (Ulm University, Germany) for technical assistance. This investigation was supported by grants awarded by the Swedish Cancer Society and The Medical Research Council. The work of SG and TH was partially supported by a grant awarded to the European ERA-NET eDEVILLI consortium.

## Materials and Methods

### Chemicals

IGEPAL CA-630 (NP40, I3021), Sodium dodecyl sulphate (SDS, L3771), N-Ethylmaleimide (NEM, E1271), Iodoacetamide (I1149), Sodium deoxycholate monohydrate (D5670), Triton X-100 (T9284), Bovine serum albumin (BSA, A7906), Tween-20 (P9416), Trizma base (Tris, 93349), Ethylenediaminetetraacetic acid disodium salt dehydrate (EDTA-E4884), Doxycycline cyclate (D9891), MG132 (M7449), Etoposide (E1383), Neocarzinostain (N9162) and Imidazole (I5513), were purchased from Sigma-Aldrich (St. Louis, MO, USA). RNase A (12091-021) and DNAzol (10503027) were purchased from Invitrogen (Carlsbad, CA, USA). Micrococcal nuclease (88216) was from Thermo Fisher Scientific (Rockford, lL, USA). Complete protease inhibitors cocktail tablets (04693116001), and phosphatase inhibitor cocktail (04906837001) were from Roche Diagnostic (Mannheim, Germany). Campthothecin was purchased from Selleckchem (Munich, Germany).

### Antibodies

The antibodies that were used in this study: mouse anti-βactin (AC-15, 1:20000) and mouse anti-FLAG (F-3165, 1:7000; IF 1:300) from Sigma-Aldrich; rabbit anti-TOP1 (A302-589A,1:5000), TOP2α (A300-054A, 1:4000), TOP2β (A300-950A, 1:2000) and 53BP1(A300-272A,1:150) were from Bethyl Laboratories (Montgomery, Texas, USA); rabbit anti phospho-histone H2A.X clone 20E3 (#9718, IF:1:100) from Cell Signaling Technology (Danvers, Massachusetts); mouse monoclonal anti-ubiquitin (P4D1, sc-8017 1:1000), mouse anti-EBV-BZLF1 (sc-53904, 1:1000) and mouse anti-BRCA1 (D-9, sc-6954, IF 1:100) from Santa Cruz Biotechnology (Dallas, Texas, USA); mouse anti-SUMO2/3 (8A2, ab81371, 1:2000) from Abcam (Cambridge, MA, USA); monoclonal rat anti-EBV-BPLF1 (1:1500) (van Gent et al., 2014) from the MAB core facility, Helmholtz Center, Munich, Germany; mouse anti-EBV-BMRF1(1:10000) and rat anti-EBV-BFRF3 (1:1000) from Dr. Jaap M. Middeldorp (VU University Medical Center, Amsterdam, Netherlands). Alexa Fluor anti-rabbit-488 (A31570, 1:1000) and anti-mouse-555 (A315721, 1:1000) conjugated secondary antibodies raised in donkey were from Thermo Fisher (Waltham, Massachusetts, USA).

### Plasmids and recombinant lentivirus vectors

Eukaryotic expression vectors encoding the N-terminal domain of the EBV large tegument protein 3xFLAG-BPLF1 (amino acid 1-235) and the catalytic mutant BPLF1-C61A(Ascherio & Munger, 2015) and the bacterial expression vector His-BPLF1(Gupta et al., 2019) were described previously. Lentiviral vectors encoding N-terminal 3xFLAG and V5 tandem tagged versions of BPLF1 aa 1-325 and the corresponding catalytic mutant BPLF1-C61A under control of the doxycycline-inducible pTight promoter were produced by cloning the corresponding open reading frames(Ascherio & Munger, 2015) into ta modified version of the pCW57.1 plasmid (gift from David Root, Addgene plasmid #41393). The Gal1/10 His6 TEV Ura S. cerevisiae expression vector (12URA-B) was a gift from Scott Gradia (Addgene plasmid #48304) a plasmid expressing human TOP2α was kindly provided by the James Berger (John Hopkins School of Medicine, Baltimore, USA). The FLAG-TOP2α construct was created by in-frame cloning the 3xFLAG coding sequence (amino acids DYKDHDGDYKDHDIDYKDDDDKL) at the N-terminus of the TOP2α open reading frame. All cloning was performed using the ligation independent cloning protocol from the QB3 Macrolab at Berkeley (macrolab.qb3.berkeley.edu). A recombinant lentivirus vector expressing the coding sequence of the EBV transactivator BZLF1 under control of a tetracycline-regulated promoter was constructed by cloning the open reading frame amplified with the primers 5’-CGACCGGTATGATGGACCCAAACTCGAC-3’ and 5’-CGACGCGTTTAGAAATTTAAGAGATCCTCGTGT-3’ into the Age I and Mlu I sites of the pTRIPZ lentiviral vector (Thermo Fisher Scientific, USA). For virus production, HEK293FT cells were co-transfected with the pTRIPZ-BZLF1, psPAX and pMD2G plasmids (Addgene, Cambridge, MA) using JetPEI (Polyplus, Illkirch, France) according to the manufacture’s protocol and cultured overnight in complete medium. After refreshing the medium, the cells were cultured for additional 48 h to allow virus production. Virus containing culture supernatant was briefly centrifuged and passed through a 0.45 μm filter to removed cell debris before aliquoting and storing at -80°C for future use.

### Cell lines and transfection

HeLa cells (ATCC RR-B51S) and HEK293T (ATCC CRL3216) cell lines were cultured in Dulbecco’s minimal essential medium (DMEM, Sigma-Aldrich), supplemented with 10% FBS (Gibco-Invitrogen) and 10 μg/ml ciprofloxacin (17850, Sigma-Aldrich) and grown in a 37°C incubator with 5% CO2. Stable HEK-rtTA-BPLF1/BPLF1-C61A cell lines were produced by lentiviral transduction followed by selection in medium containing 2μg/puromycin for 2 weeks. Expression of FLAG-BPLF1/BPLF1-C61A was induced by treatment with 1.5 μg/ml doxycycline and confirmed by anti-FLAG immunofluorescence and Western blot analysis. Clones expressing high levels of the transduced proteins were selected by limiting dilution. HeLa cells were transiently transfected with plasmids expressing FLAG-tagged version of BPLF1/BPLF1-C61A using the lipofectamine 2000 (Invitrogen, California, USA) or jetPEI® (Polyplus transfection, Illkirch FR) DNA transfection reagent according to the protocols recommended by the manufacturer.

### Production of EBV immortalized lymphoblastoid cell lines (LCLs)

Peripheral blood mononuclear cells were purified from Buffy coats (Blood Bank, Karolinska University Hospital, Stockholm, Sweden) by Ficoll-Paque (Lymphoprep, Axis-shield PoC AS, Oslo, Norway) density gradient centrifugation, and B-cells were affinity-purified using CD19 microbeads (MACS MicroBeads, Miltenyi Biotec, Bergisch Gladbach, Germany) resulting in >95% pure B-cell populations. Infectious EBV encoding wild type or catalytic mutant BPLF1 were rescued for HEK293-EBV cells as previously described(Gupta et al., 2018). One million B-cells were incubated in 1 ml virus preparation for 1.5 h at 37°C, followed by the addition of fresh complete medium and incubation at 37°C in a 5% CO2 incubator until immortalized LCLs were established. Sublines expressing a doxycycline-inducible BZLF1 transactivator were produced by culturing 10^6^ LCL cells with the recombinant lentivirus in presence of 8 μg/ml polybrene (TR-1003-G, Sigma-Aldrich) for 24 hours followed by replacement of the infection medium with fresh complete medium. The transduced cells were selected in medium containing 0.8μg/ml (LCL-BPLF1) or 0.25μg/ml (LCL-BPLF1-C61A) puromycin for one or two weeks.

### Immunofluorescence

Transfected HeLa and HEK-Tta-BPLF1/BPLF1-C61A cells were grown on coverslips and induced with 1.5 μg/ml doxycycline for 24 h. For immunofluorescence analysis, the cells were fixed with 4% formaldehyde for 20 min, followed by permeabilization with 0.05% Triton X-100 in PBS for 5 min and blocking in PBS containing 4% bovine serum albumin for 40 min. After incubation for 1 h with primary antibodies and washing 3×5 min in PBS, the cells were incubated for 1 h with the appropriate Alexa Fluor-conjugated secondary antibodies, followed by washing and mounting in Vectashield-containing DAPI (Vector Laboratories, Inc. Burlingame, CA, USA). Images were acquired using a fluorescence microscope (Leica DM RA2, Leica Microsystems, Wetzlar, Germany) equipped with a CCD camera (C4742-95, Hamamatsu, Japan). Fluorescence intensity was quantified using the ImageJ® software.

### Western blots

Cells were lysed in RIPA buffer (25 mM Tris•HCl pH 7.6, 150 mM NaCl, 1% Igepal, 1% sodium deoxycholate, 2% SDS) supplemented with protease inhibitor cocktail. Loading buffer (Invitrogen) was added to each sample followed by boiling for 10 min at 100°C. The lysates were fractionated in acrylamide Bis-Tris 4-12% gradient gel (Life Technologies Corporation, Carlsbad, USA). After transfer to PVDF membranes (Millipore Corporation, Billerica, MA, USA), the blots were blocked in TBS (VWR, Radnor, Pennsylvania, USA) containing 0.1% Tween-20 and 5% non-fat milk, and the membranes were incubated with the primary antibodies diluted in blocking buffer for 1 h at room temperature or over-night at 4°C followed by washing and incubation for 1 h with the appropriate horseradish peroxidase-conjugated secondary antibodies. The immunocomplexes were visualized by enhanced chemiluminescence (GE Healthcare AB, Uppsala, SE). For detecting topoisomerase-DNA adducts after treatment with topoisomerase poisons, the cells were lysed in alkaline buffer(Ban et al., 2013). Briefly, cells treated for the indicated time with 5 μM Campthothecine or 80 μM Etoposide were lysed in 100 μl in buffer containing 200 mM NaOH, 2 mM EDTA, followed by the addition of 100 μl of 1M HEPES buffer (pH 7.4). Nucleic acids were removed by addition of 10 μl 100 mM CaCl2, 2 μl 1M DTT, and 200U of micrococcal nuclease followed by incubation at 37°C for 20 min. Seventy μl of 4xLDS loading buffer (Invitrogen) were added to each sample followed by boiling for 10 min at 100°C before SDS-PAGE fractionation and western blot analysis.

### Immunoprecipitation and pull-down assays

Cells were harvested 48h after transfection and lysed in NP40 lysis buffer (150 mM NaCl, 50 mM Tris-HCl pH7.6, 5mM MgCl2, 1mM EDTA, 1% Igepal, 1 mM DTT, 10% glycerol) supplemented with protease/phosphatase inhibitor cocktail, 20 mM NEM and 20 mM Iodoacetamide for 30 min on ice. For immunoprecipitations under denaturing condition the lysis buffer was supplemented with 1% SDS followed by dilution to 0.1% SDS. For BPLF1/BPLF1-C61A co-immunoprecipitation, the lysates were incubated for 3 h with 50 μl anti-FLAG packed agarose affinity gel (A-2220; Sigma) at 4 °C with rotation. After washing 4 times with lysis buffer, the immunocomplexes were eluted with FLAG peptide (F4799; Sigma). For TOP2α and TOP2β immunoprecipitation, specific antibodies were added to cell lysates and incubated at 4°C for 3 h with rotation. The protein-antibody complexes were captured with protein-G coupled Sepharose beads (GE Healthcare) by incubation at 4 °C for 1 h. The beads were washed 4 times with lysis buffer followed by boiling in 2xSDS-PAGE loading for 10 min at 100°C. The production of 6xHis-BPLF1 in bacteria and purification of the recombinant protein were done as previously described(Gupta et al., 2019). Recombinant human TOP2α was expressed and purified according to a previously published protocol with slight modifications(Lee et al., 2017). Briefly, URA-deficient yeast (kindly provided by Lena Ström, CMB Karolinska Institutet) were transformed with the TOP2α expression plasmid, grown initially in uracil-deficient media, then in YPLG (1% yeast extract, 2% peptone, 2% sodium DL-lactate, 1.5% glycerol) before induction of expression by addition of 2% galactose. The yeast was harvested by centrifugation and snap-frozen in liquid nitrogen. Proteins were extracted using a cryo-mill, and the filtered lysate was passed sequentially through HisTrap Excel nickel and HiTrap CP cation exchange columns (GE Healthcare) to purify the tagged TOP2α protein before incubation overnight with His-tagged TEV protease. The following day, the protein was passed through a HisTrap column to remove the cleaved His-tag and the TEV protease. The TOP2α protein was further purified on a Superdex 200 16/60 column, concentrated, and stored at -80°C. Relaxation and decatenation assays along with western blotting were performed to confirm protein purity and activity. Equimolar concentration of purified His-BPLF1 (0.35 μg) and FLAG-TOP2α (2 μg) were incubated in binding buffer (100 mM NaCl, 50 mM Tris-HCl, 1mM DTT, 0.5% Igepal) for 20 min at 4°C. Anti-FLAG agarose affinity gel (A-2220; Sigma) or Ni-NTA beads (Qiagen) were added followed by incubation for 60 min or 20 min at 4°C with rotation. The beads were intensively washed, and bound proteins were eluted with FLAG peptide or 300 mM imidazole in buffer containing 50 mM Tris-HCl pH 7.6, 50 mM NaCl and 1 mM DTT.

### Rapid approach to DNA adduct recovery (RADAR) assay

TOP2ccs were isolated by RADAR assays as described(Kiianitsa & Maizels, 2013). Briefly, cells cultured in 6 well plates were treated with 80 μM etoposide for 30 min or 4 h and then lysed in 800 μl DNAzol. Following the addition of 400 μl absolute ethanol, the lysates were cooled at -20°C and then centrifuged at 14000 rpm for 20 min at 4°C. After repeated washing in 75% ethanol the nucleic acid pellets were dissolved in 100 μl H_2_O at 37°C for 15 minutes, followed by treatment with 100 μg/ml RNaseA. The concentration of DNA was measured and 10 μg DNA from each sample were treated with 250 U micrococcal nuclease supplemented with 5 mM CaCl_2_ before the addition of loading buffer and detection of trapped protein by western blot.

### Reverse transcription and real-time PCR

Total RNA was isolated using the Quick-RNA MiniPrep kit (Zymo Research, Irvine, CA, USA) with in-column DNase treatment according to the instructions of the manufacturer. One microgram of total RNA was reverse transcribed using SuperScript VILO cDNA Synthesis kit (Invitrogen). PCR amplification was performed with the LC FastStart DNA master SYBR green I kit in a LightCycler 1.2 instrument (Roche Diagnostic) using the following specific primers: TOP1 5’-AGTGGAAAGAAGTCCGGCATGA-3’, 5’-GCCAGTCCTTCTCACCCT TGAT-3’; TOP2α 5’-AAGCCCAGCAAAAGGTTCCA-3’, 5’-TGGCTTCAACAGCCTCCA AT-3’; TOP2β 5’-GGTTCGTGTAGAGGGGTCAA-3’, 5’-CCCAGTTTCATCCAATTTGT C-3’; BPLF1 5’-CATACACCGTGCGAAAAGAA-3’, 5’-GATGGCGGGTAATACATGCT-3’; and MLN51 (Metastatic Lymph Node 51) 5’-CAAGGAAGGTCGTGCTGGTT-3’, 5-AC CAGACCGGCCACCAT-3’as endogenous control gene. The PCR reactions were denatured at 95°C for 10 min, followed by 40 cycles at 95°C for 8 sec, 60°C for 5 sed, 72°C for 8 sec. The relative levels of mRNA were determined from the standard curve using MLN51 as reference.

### MTT assay

For assay of cell viability, 2×10^4^ HEK-rtTABPLF1/BPLF1-C61A or 5×10^4^ LCLEBV-BPLF1/BPLF1-C61A were plated in 150 μl medium in triplicate wells of a 96 well plate without or with the addition of the indicated concentrations of Etoposide. After incubation for 20 h at 37°C in a 5% CO2 incubator, 50 μl culture medium containing 1 mg/ml Methylthiazolyldiphenyl-tetrazolium bromide (MTT, M5655, Sigma-Aldrich) were added to the wells followed by incubation for additional 4 h. The MTT formazan crystals produced by mitochondrial dehydrogenases in living cells were solubilized by the addition of 50 μl 10% SDS and O.D. was measured at 540 nm in a plate reader. Relative viability was calculated after subtraction of the background O.D. of media alone.

### EBV DNA replication and release of infectious virus

Virus replication and the release of infectious virus were monitored after induction with 1.5 μg/ml Doxycycline for 3 days in cell pellets and culture supernatants by quantitative PCR. Briefly, DNA was isolated from cell pellets and culture supernatants cleared of cell debris by centrifugation of 5 min at 14000 rpm and treated with 20 U/ml DNase I (Promega, Madison, WI, USA) to remove free viral DNA, using the DNeasy Blood & Tissue Kit (Qiagen, Hilden, Germany). Quantitative PCR was performed Quantitative PCR was performed as described above with primers specific for a unique sequence in EBNA1 5’-GGCAGTGGACCTCAAAG AAG-3’, 5’-CTATGTCTTGGCCCTGATCC-3’ and the cellular EF1α Elongation factor α 5’-CTGAACCATCCAGGCCAAAT-3’, 5’-GCCGTGTGGCAATCCAAT-3’ as reference. Virus replication was calculated as the ratio between the amount of viral DNA in induced versus untreated cells.

