## Supplementary figures for "The Epstein-Barr virus ubiquitin deconjugase BPLF1 regulates the activity of Topoisomerase II during virus replication"

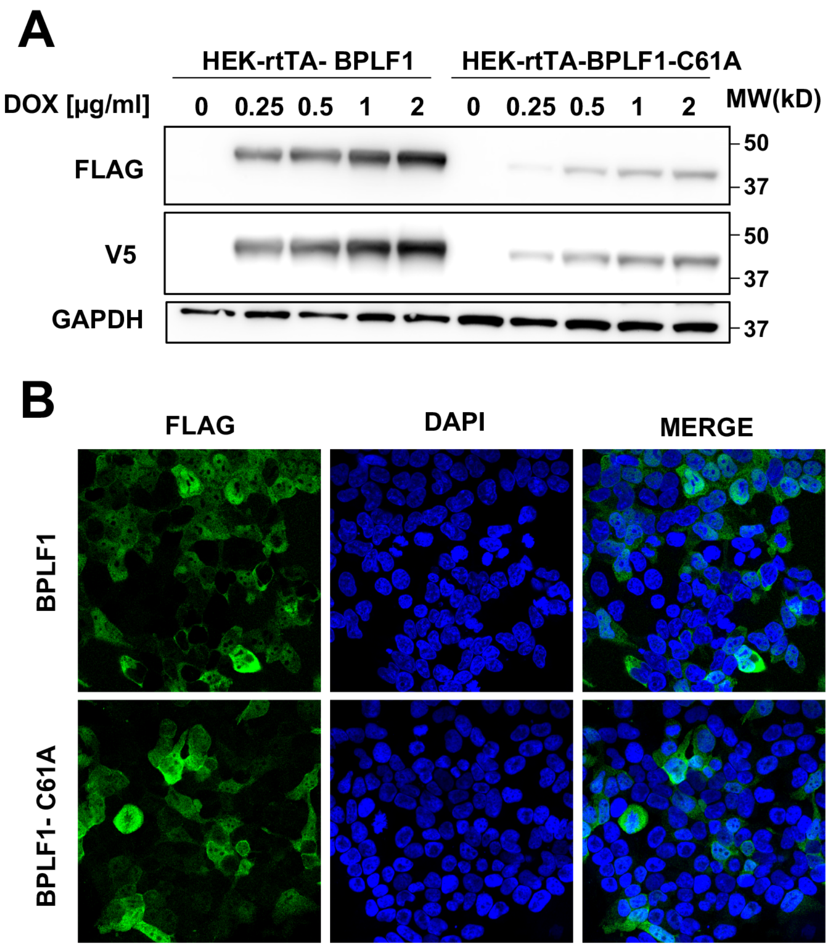


**Figure S1. Characterization of the HEK-rtTA-BPLF1/BPLF1-C61A cell lines. A)** Expression of BPLF1/BPLF1-C61A was detected in western blots of cells treated for 24 h with the indicated amount of Dox using antibodies to the FLAG tag. Lower steady-state levels of BPLF1-C61A were usually detected due to rapid protein turnover. **B)** Representative micrographs illustrating the expression of BPLF1/BPLF1C61A in untreated and Dox-treated cells. Confocal images were obtained at 40x lens objective magnification. BPLF1 is in green and cell nuclei were stained with DAPI (blue). Strong FLAG fluorescence was detected in approximately 50% of the induced cells.

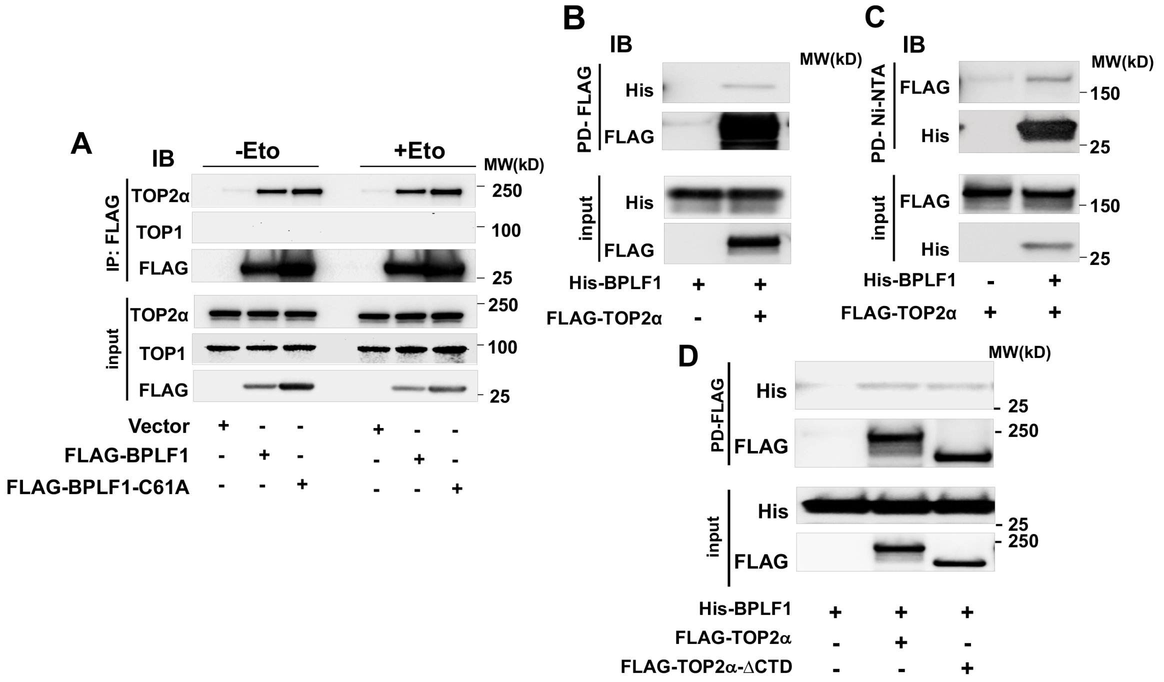


**Figure S2. BPLF1 does not interact with TOP1 and recombinant BPLF1 binds to TOP2.** **(A)** HEK293T cells were transfected with FLAG-BPLF1, FLAG-BPLF1-C61A, or empty FLAG-vector and then treated with 40 μM Etoposide for 30 min. Cell lysates were immunoprecipitated with anti-FLAG conjugated agarose beads and western blots were probed with the indicated antibodies. TOP2α was readily detected in the immunoprecipitates while TOP1 was consistently absent. Representative western blots from one of two independents experiments giving similar results are shown. (**B,C,D**) The interaction of yeast expressed FLAG-TOP2α or TOP2α lacking the C-terminal domain (FLAG-TOP2α-ΔCTD) with bacterially expressed His-BPLF1 was assayed in pull-down assays. Equimolar amounts of the proteins were mixed and FLAG **(B, D)** or Ni-NTA (**C**) pull-downs were probed with antibodies specific for FLAG or BPLF1. A weak interaction of BPLF1 with TOP2α was detected independently of the presence of the TOP2α C-terminal domain. Western blots from one representative experiment out of two are shown in the figure.

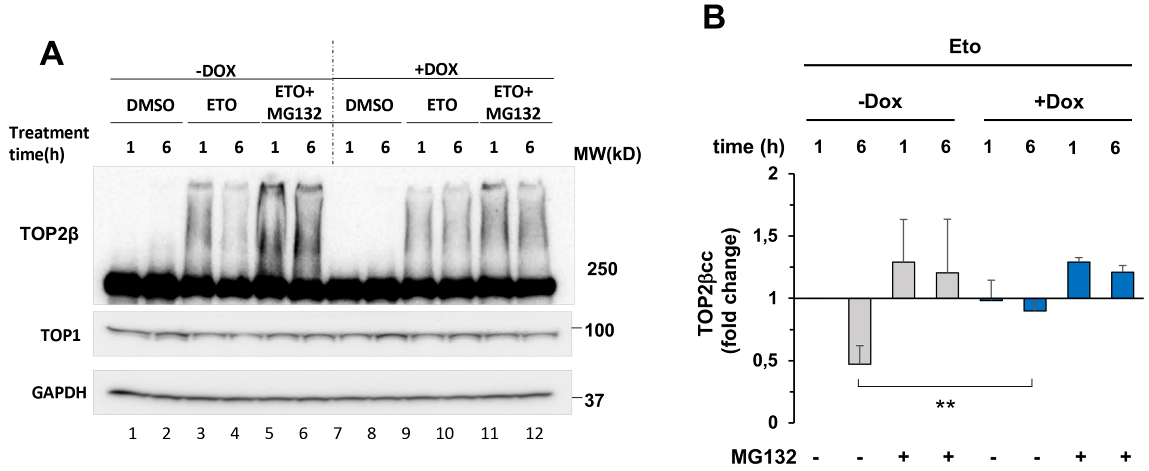


**Figure S3. BPLF1 inhibits the resolution of TOP2cc. (A)** HEK-rtTA-BPLF1 cells were cultured with or without 1.5μg/ml Dox for 24 h and then treated with 80 μM Etoposide alone or together with 10 μM MG132. Cells harvested after 1 h or 6 h were lysed in alkaline buffer and the formation of TOP2cc was investigated by probing western blots with the TOP2β antibody. The TOP2cc are visualized as smears of DNA cross-linked TOP2β above the main band. Probing with the anti-TOP1 antibody confirmed the selective induction of TOP2βcc in Etoposide treated cells. GAPDH was used as the loading control. Western blots from one representative of three independent experiments are shown in the figure. (**B**) Densitometry quantification confirming the stabilization and TOP2βcc in BPLF1 expressing cells. The mean ± SD of two independent experiments is shown. **P<0.01

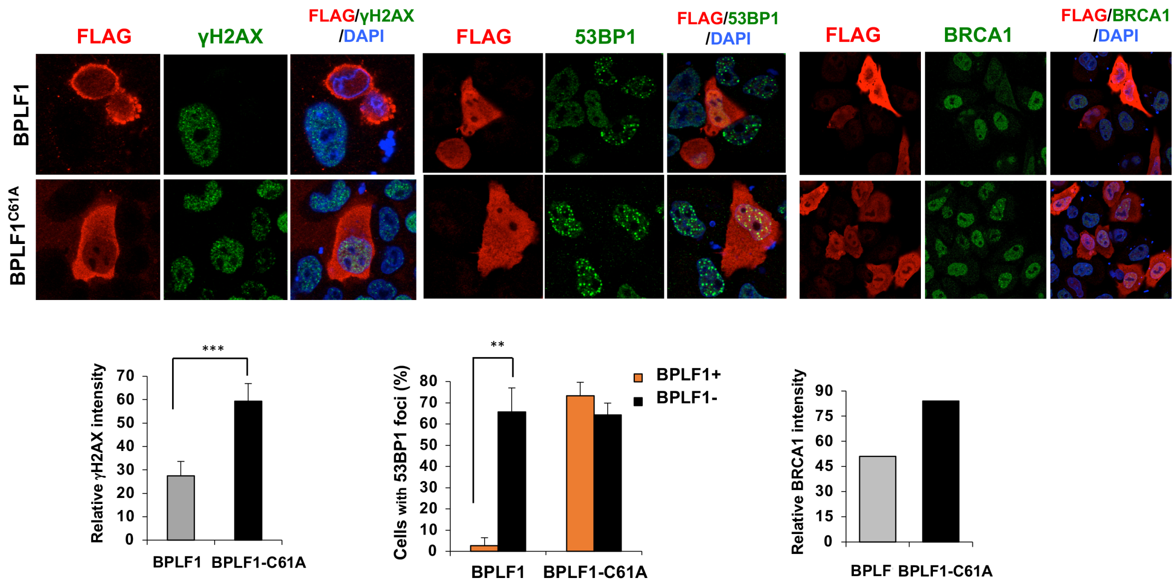


**Figure S4. Transfection of catalytically active BPLF1 inhibits activation of the DDR and DNA repair in etoposide treated HeLa cells**. HeLa cells transiently transfected with plasmids expressing FLAG-BPLF1/BPLF1-C61A were treated for 6 h with 40 μM etoposide before fixation and staining with the indicated antibodies. Representative micrographs of cells co-stained with antibodies to FLAG, the DNA-DSB maker γH2AX, and the DNA repair markers 53BP1 and BRCA1. Expression of catalytically active BPLF1 was associated with decrease γH2AX and BRCA1 fluorescence and failure to accumulate 53BP1 foci. Images from one representative experiment out of three are shown. The intensity of γH2AX and BRCA1 fluorescence and the number of cells showing ≥ 2 53BP1 foci were quantified in BPLF1 positive and negative cells from the same transfection experiment using the ImageJ software. Mean ± SD of two or three independent experiments where a minimum of 50 BPLF1 positive and 50 BPLF1 negative cells was scored in each condition.

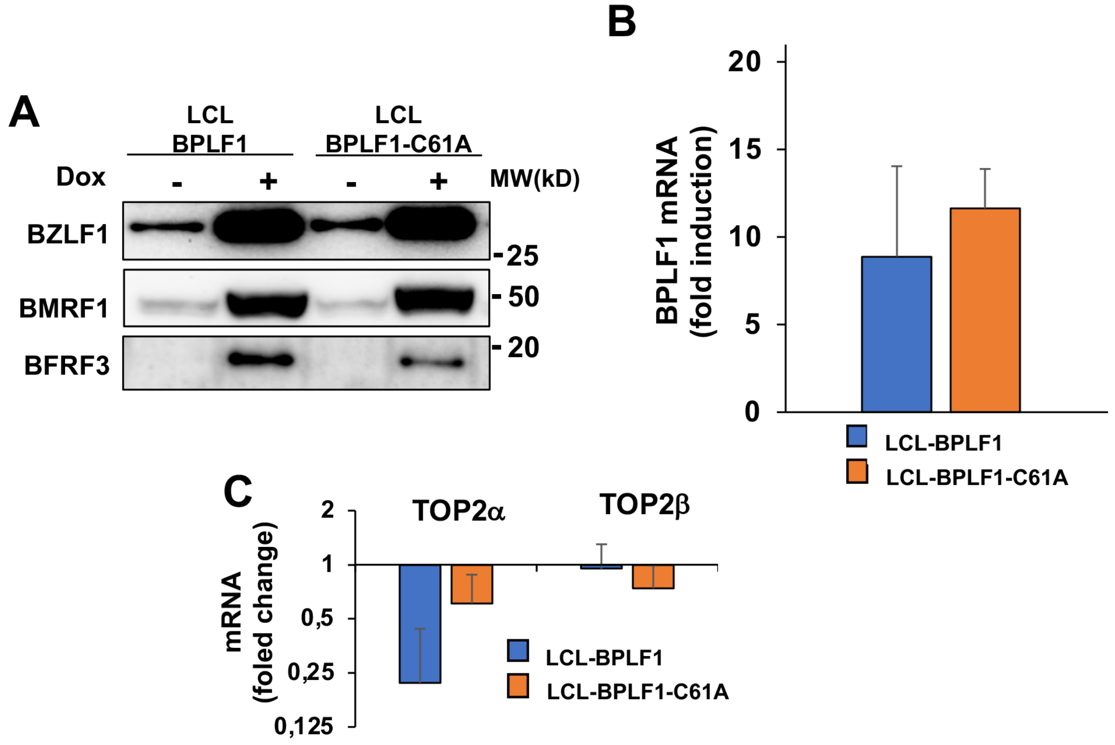


**Figure S5. Induction of the productive virus cycle and quantification of TOP2 mRNA in LCLs carrying recombinant EBV expressing wild type and mutant BPLF1.** The productive cycle was induced in LCL-EBV-BPLF1/BPLF1-C61A by culture for 72 h in the presence of 1.5 µg/ml Dox. **(A)** Viral gene expression was assessed by probing western blots of total cell lysates with the indicated antibodies to the immediate early antigen BZLF1, the early antigen BMRF1 and the late antigen BFRF3. The expression of BPLF1 **(B)**, TOP2α and TOP2β **(C)** mRNA was quantified by qPCR. The mean ± SD fold increase relative to uninduced controls recorded in three independent experiments is shown in the figure.
